# Cell-Type Specific Responses to Associative Learning in the Primary Motor Cortex

**DOI:** 10.1101/2021.08.08.455571

**Authors:** Candice Lee, Emerson Harkin, Richard Naud, Simon X. Chen

## Abstract

The primary motor cortex (M1) is known to be a critical site for movement initiation and motor learning. Surprisingly, it has also been shown to possess reward-related activity, presumably to facilitate reward-based learning of new movements. However, whether reward-related signals are represented among different cell types in M1, and whether their response properties change after cue-reward conditioning remains unclear. Here, we performed longitudinal *in vivo* two-photon Ca^2+^ imaging to monitor the activity of different neuronal cell types in M1 while mice engaged in a classical conditioning task. Our results demonstrate that most of the major neuronal cell types in M1 showed robust but differential responses to both cue and reward stimuli, and their response properties undergo cell-type specific modifications after associative learning. PV-INs’ responses became more reliable to the cue stimulus, while VIP-INs’ responses became more reliable to the reward stimulus. PNs only showed robust response to the novel reward stimulus, and they habituated to it after associative learning. Lastly, SOM-IN responses emerged and became more reliable to both conditioned cue and reward stimuli after conditioning. These observations suggest that cue- and reward-related signals are represented among different neuronal cell types in M1, and the distinct modifications they undergo during associative learning could be essential in triggering different aspects of local circuit reorganization in M1 during reward-based motor skill learning.

## INTRODUCTION

The primary motor cortex (M1) is an essential site for movement execution and motor learning. Within M1, neurons encode movement goals and movement kinematics (Georgopoulos, Ashe, Smyrnis, & Taira, 1992; Moran & Schwartz, 1999; Peters, Chen, & Komiyama, 2014). Intriguingly, neurons in M1 have also been reported to show reward-related activity. *In vivo* recording studies performed in nonhuman primates found neurons in M1 that encode reward anticipation, reward delivery, and mismatches between reward delivery and reward expectation (Marsh et al., 2015; Ramakrishnan et al., 2017; Ramkumar et al., 2016). In human subjects, reward has also been shown to modulate M1 activity, likely through an inhibitory circuit dependent mechanism (Thabit et al., 2011). However, it remains unclear how reward-related responses are represented in M1, and if the representation changes with associative learning.

It was recently shown that in well-trained mice performing a skilled reaching task, a subset of layer 2/3 (L2/3) pyramidal neurons (PNs) in M1 specifically report successful, but not failed, reach-and-grasp movements. In contrast, a different subset of PNs report only failed reach-and-grasp movements (Levy et al., 2020). Since the ability to use past experience to learn action-outcome associations is critical to survival, encoding the outcome in M1 may be an important part of motor skill learning. It is widely accepted that associative learning using reinforcement can accelerate and enhance learning (Abe et al., 2011; Nikooyan & Ahmed, 2015). In the case of motor learning, studies have demonstrated that positive feedback (reward) facilitates motor memory retention and negative feedback (punishment) speeds up the learning process (Galea et al., 2015). One hypothesis is that during learning, reward signals in the brain, together with neuromodulators and synaptic plasticity, are involved in potentiating and optimizing the neural circuitry in M1 that underlies the rewarded movement. Implementing such a learning process would necessitate the interplay between different cell types within the local microcircuitry (Richards et al., 2019).

M1, like other cortical areas, is densely packed with PNs and diverse inhibitory interneuron (IN) types and is wired in a delicately balanced and intricate circuit. Different IN subtypes have been shown to have distinct gene expression profiles, electrophysiological properties, and connectivity motifs (Fishell & Rudy, 2011; Markram et al., 2004). Somatostatin-, parvalbumin-, and vasoactive intestinal peptide-expressing inhibitory neurons (SOM-INs, PV-INs and VIP-INs, respectively) are three major non-overlapping subtypes of GABAergic neurons that broadly form a common microcircuit motif in the cortex. Some studies have demonstrated that SOM-INs preferentially target distal dendrites of PNs to filter synaptic inputs, fast-spiking PV-INs preferentially target perisomatic regions of PNs enabling strong inhibition of spiking, and VIP-INs regulate local microcircuits by controlling other local INs (Pfeffer et al., 2013). Due to their diverse properties and strategic connectivity motifs, these INs exert fine control over local network activity and provide a potential mechanism for how the brain processes reward signals and ultimately uses this information to optimize neural activity related to learned motor skills.

Multiple studies using *in vivo* opto-recording in the primary visual cortex have shown that visual orientation selectivity in PNs is modulated and sharpened by PV- and SOM-INs (Atallah et al., 2012; Lee et al., 2012; Wilson et al., 2012). In the primary auditory cortex, PV- and SOM-INs exert analogous control over PN frequency tuning (Seybold et al., 2015). Moreover, in the auditory cortex, prefrontal cortex, and basolateral amygdala, reinforcement signals such as reward and punishment have been shown to recruit VIP-INs, which in turn, inhibit SOM- and PV-INs (Krabbe et al., 2019; Pi et al., 2013). The subtype-specific roles of these INs have long been elusive, but a complex picture is emerging where INs are not only responsible for maintaining a delicate balance of excitation and inhibition, but are also actively involved in processing activity in cortex (Lee et al., 2020; Wood et al., 2017).

Here, we employed chronic *in vivo* two-photon imaging, combined with a head-fixed classical conditioning task, to monitor the activity of the same population of PNs, PV-INs, SOM-INs, or VIP-INs before and after associative learning to investigate whether and how conditioned cue and unconditioned reward signals are represented among different neuronal cell types in M1. Our results demonstrate that all four major cell types in M1 show distinct responses to conditioned cue and reward stimuli, and their response properties undergo cell-type specific modifications after associative learning. Notably, PV-INs and VIP-INs exhibited stimulus-specific modifications, in which PV-INs became more reliably responsive to the cue but not to the reward stimulus, whereas VIP-INs became more reliable to the reward but not to the cue stimulus. PNs initially showed robust responses to novel reward stimuli but habituated to it after associative learning. Lastly, SOM-IN responses emerged and became more reliable to both conditioned cue and reward stimuli following associative learning. Taken together, these results show that cue- and reward-related signals are represented among major neuronal cell types in M1, and they undergo cell-type specific modifications during associative learning, indicating they may have distinct roles in integrating reinforcement signals to promote circuit reorganization in M1 during motor skill learning.

## RESULTS

To understand how reward-associated signals are represented within the local microcircuitry in M1 before and after associative learning, we established a head-fixed auditory cued reward conditioning task, which allowed us to combine the task with *in vivo* two-photon Ca^2+^ imaging to examine the response properties of different neuronal cell type populations in awake and behaving mice (**Figure 1A**). In this task, water-restricted mice were exposed to a conditioned stimulus (CS; auditory tone, 1s duration), followed by a 1.5s delay and then the delivery of the unconditioned stimulus (US; water reward, ∼10 µL). Mice were trained for ∼30-35 trials/session (1 session/day for 7 days) with a randomly varied inter-trial interval (ITI) between 60 – 120s (**Figure 1B**). Since M1 is known to be involved in movement initiation and motor skill learning, we chose to use a simple classical conditioning task with just an auditory tone paired with reward and omitted any additional training where mice would be required to learn a new movement. The rationale for this is that many neuronal cell types, including PNs, PV- and SOM-INs, have been shown to undergo modifications when mice acquire new movements (Chen et al., 2015; Cichon & Gan, 2015; Donato et al., 2013; Xu et al., 2009). However, since licking is an innate movement that does not induce plastic changes in adult mice (Chen et al., 2015; Komiyama et al., 2010; Peters et al., 2014), we can reliably attribute changes in neuronal activity over the course of the task to associative learning, rather than to motor learning.

**Figure 1.**
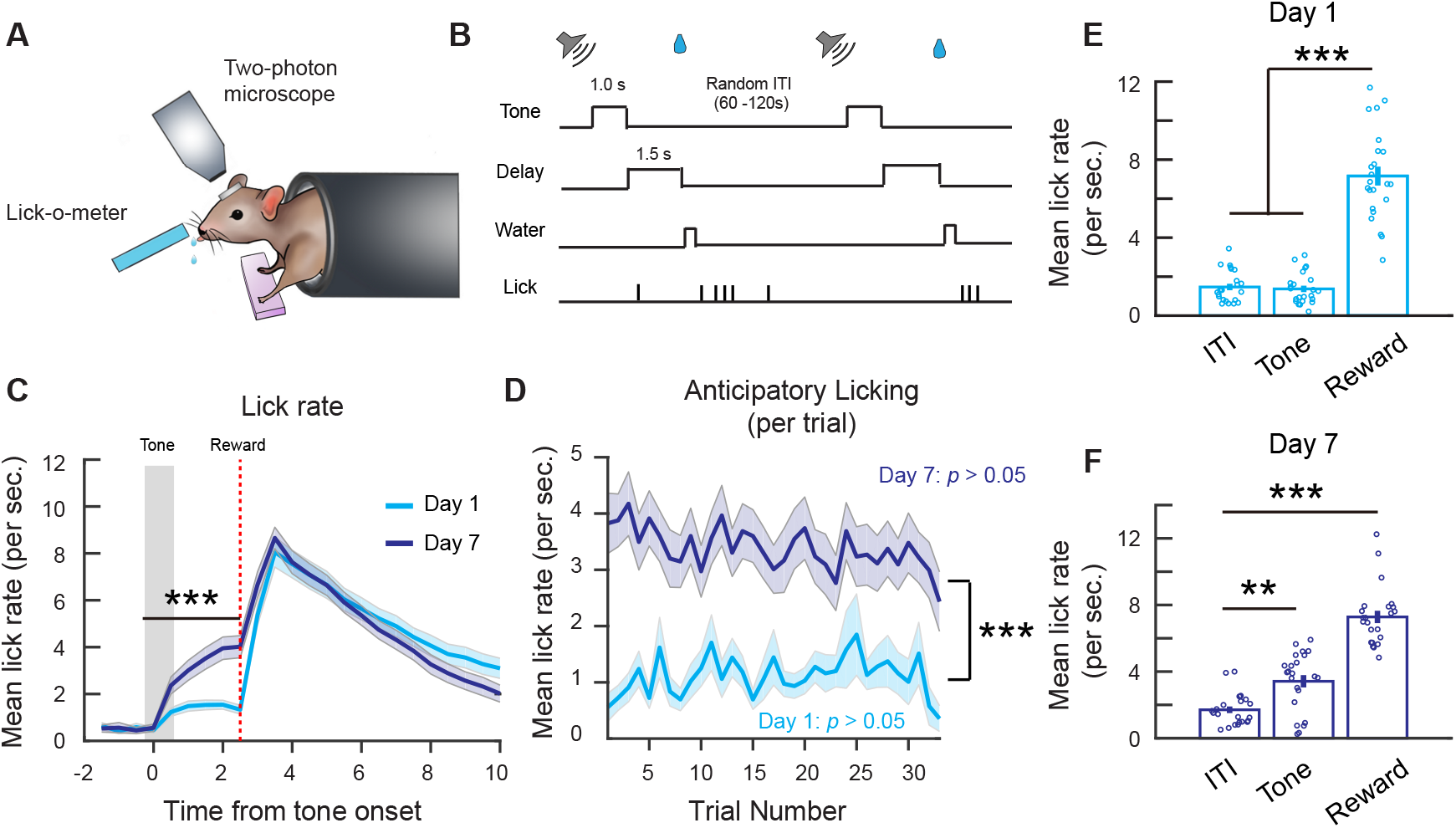
Associative learning during a head-fixed classical conditioning task. **(A)** Schematic of head-fixed classical conditioning task. **(B)** Trial structure. **(C)** Mean lick rate per second on day 1 and day 7. Binned over 0.5s intervals. Lick rate following tone onset up to reward delivery time is higher on day 7. Two-way ANOVA, *** *p < 1*×*10*^*-3*^, effect of time: *p < 1*×*10*^*-3*^, effect of day: *p < 1*×*10*^*-3*^. **(D)** Mean anticipatory lick rate across trials within day 1 and day 7 sessions. Mean anticipatory lick rate was calculated from tone onset to end of delay period. Two-way ANOVA, effect of trial number: *p < 1*×*10*^*-3*^, effect of day: *p < 1*×*10*^*-3*^. **(E)** Mean lick rate during the first 2.5s of ITI lick bouts, 2.5s following tone onset and 2.5s following reward delivery on day 1. One-way ANOVA with Tukey-Kramer correction for multiple comparisons, ITI vs. tone: *p = 0*.*97*, ITI vs. reward: *p < 1*×*10*^*-3*^, tone vs. reward: *p < 1*×*10*^*-3*^. **(F)** Mean lick rate during the first 2.5s of ITI lick bouts, 2.5s following tone onset and 2.5s following reward delivery on day 7. One-way ANOVA with Tukey-Kramer correction for multiple comparisons. ITI vs. tone: *p = 1*.*08* × *10*^*-3*^, ITI vs. reward: *p < 1*×*10*^*-3*^. n = 23 mice. Error bars show SEM.

Mice learned to associate the CS with the reward after 7 days, shown by an increase in anticipatory lick rate, a conditioned response, following the tone onset on day 7 compared to day 1 (**Figure 1C**). On a trial-by-trial basis, anticipatory lick rate did not change significantly within a single session on both day 1 and day 7, implying limited within-session improvements (**Figure 1D**). To ensure the increase in lick rate was specific to the CS, we compared the mean lick rate during the CS and reward period to the lick rate during self-initiated spontaneous lick bouts in the ITI (in the absence of the CS or reward). To be consistent with the 2.5s analysis window for CS responses, we analysed the first 2.5s of ITI lick bouts and 2.5s following reward delivery. On day 1, the mean lick rate during ITI lick bouts (1.47 ± 0.17/s) and the CS (1.37 ± 0.17/s) were similar, while the lick rate following reward delivery was significantly higher (7.16 ± 0.48/s). In contrast, on day 7 following associative learning, the lick rate during the CS period (3.42 ± 0.37/s) was significantly higher than during ITI lick bouts (1.71 ± 0.2/s), demonstrating the mice effectively learned the CS-reward association by day 7 (**Figure 1E, F**).

To investigate the activity of different neuronal cell types during this task, we used *in vivo* two-photon Ca^2+^ imaging of different cell type populations. To target PNs in M1, we injected an adeno-associated virus (AAV) carrying a Ca^2+^ indicator (GCaMP6f) driven by the CaMKII promoter (AAV1.CaMKIIa.GCaMP6f) into M1 of wildtype B6129S mice (**Figure 2A**). After 3-5 weeks, we recorded the activity of hundreds of L2/3 PNs using two-photon microscopy in awake mice while they underwent the head-fixed conditioning task, and we tracked the same population of neurons on day 1 and day 7. We identified all the active neurons within a session, irrespective of the behavioural task (see Methods), and sorted neurons by the timing of their peak activity relative to the CS onset. It was apparent that there are subpopulations of neurons more responsive to CS, reward, or both (**Figure 2B, C**). We also repeated the experiments to examine if the major IN subtypes in M1 also respond to the CS and reward during the conditioning task. To do this, we injected AAV-Syn-Flex-GCaMP6f in PV-Cre, SOM-Cre or VIP-Cre transgenic mice to selectively express GCaMP6f in PV-INs, SOM-INs, or VIP-INs, respectively, and then performed *in vivo* two-photon Ca^2+^ imaging to monitor the response properties of the same population of INs on day 1 and day 7 after associating learning. We also compared the mean percentage of active cells within the entire session and confirmed all cell types had a similar proportion of active cells (irrespective of the behavioural task) on day 1 and day 7 (**Figure 2D**).

**Figure 2.**
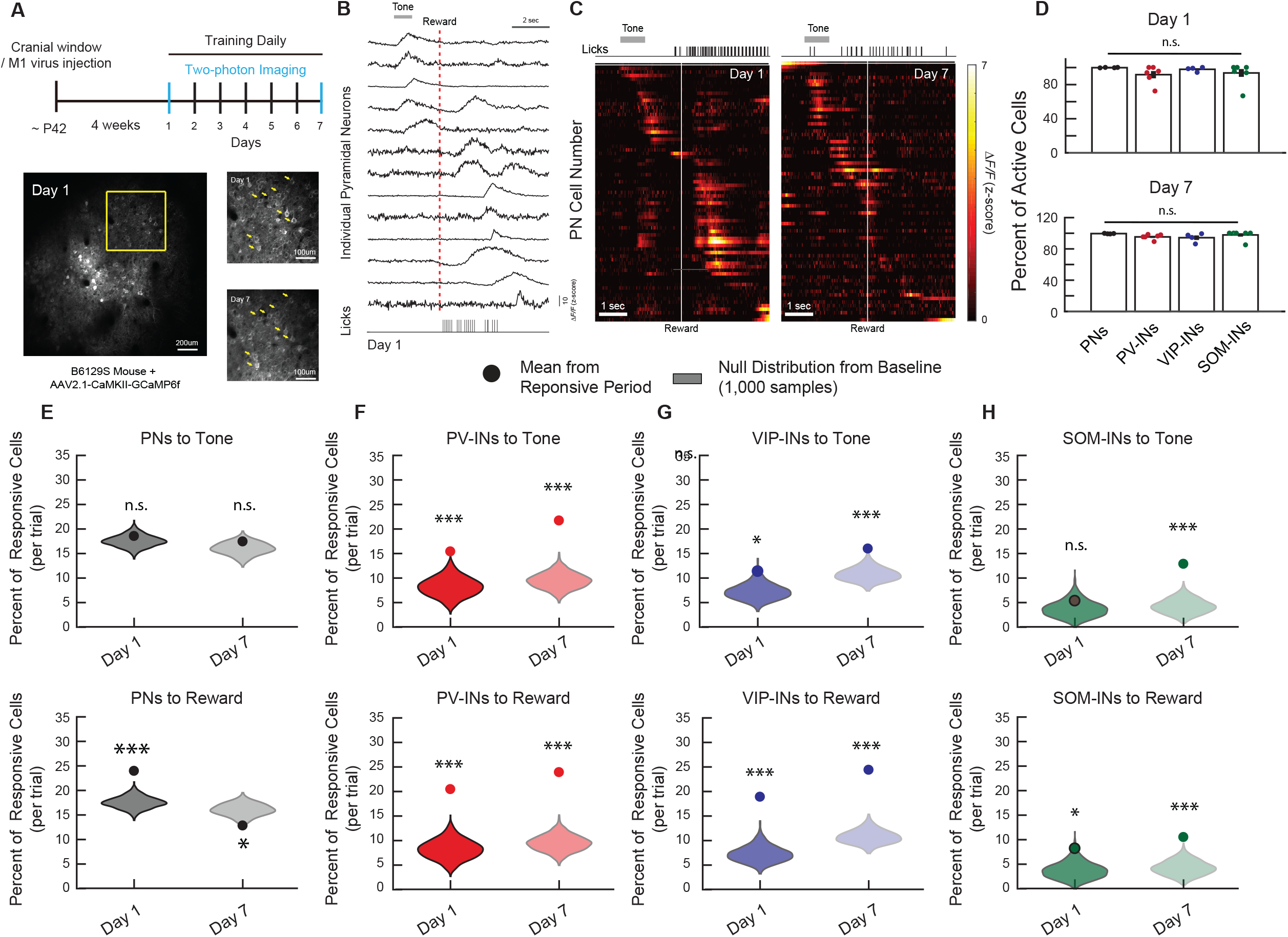
Longitudinal *in vivo* Ca^2+^ imaging of neuronal responses in M1 during a classical conditioning task. **(A)** Experimental timeline (top). *In vivo* two-photon imaging of L2/3 PNs expressing GCaMP6f in M1 (bottom left). The same population can be tracked from day 1 to day 7 (bottom right). Yellow arrows indicate example tracked neurons across days. **(B)** Z-scored fluorescence traces from 13 neurons (top), and the corresponding licking measured with the lick-o-meter (bottom) from the same mouse and same trial on day 1. Grey bar represents the timing of the auditory cue stimuli. Dotted red line indicates the onset of water reward delivery. **(C)** Z-scored activity of all the active neurons from an example mouse during one representative trial on day 1 and day 7, sorted by timing of maximum activity following the tone onset. Grey bar represents the timing of the tone stimulus. White line indicates the onset of water reward delivery. **(D)** Mean percent of active neurons within a session, irrespective of the behavioural task, for PNs, PV-INs, VIP-INs and SOM-INs on day 1 and day 7. All cell types showed a similar percentage of active neurons. One-way ANOVA, n.s., non-significant, Day 1: *p = 0*.*38*, Day 7: *p = 0*.*22*. Error bars show SEM. **(E-H)** Mean percent of responsive neurons to tone (top) and reward (bottom) within 2.5s of tone/reward stimulus onset for each cell type. Violin plots show null distribution of percentage of responsive neurons made by re-sampling each mouse and shuffling the session 1,000 times (see Methods). The circle represents the mean percentage of tone- or reward-responsive neurons. Monte Carlo with Bonferroni correction, n.s., non-significant, * *p < 0*.*05*, *** *p < 1*×*10*^*-3*^. PN Tone Day 1: *p = 0*.*582*, Tone Day 7: *p = 0*.*423*, Reward Day 1: *p < 1*×*10*^*-3*^, Reward Day 7: *p = 0*.*015*, n = 1029 cells from 6 mice **(E)**, PV-IN Tone Day 1 *p < 1*×*10*^*-3*^, Tone Day 7: *p < 1*×*10*^*-3*^, Reward Day 1: *p < 1*×*10*^*-3*^, Reward Day 7: *p < 1*×*10*^*-3*^, n = 316 cells from 6 mice **(F)**, VIP-IN Tone Day 1: *p* = *0*.*039*, Tone Day 7: *p < 1*×*10*^*-3*^, Reward Day 1: *p < 1*×*10*^*-3*^, Reward Day 7: *p < 1*×*10*^*-3*^, n = 316 cells from 6 mice n = 407 cells from 4 mice **(G)**, SOM-IN Tone Day 1: *p* = 0.47, Tone Day 7: *p < 1*×*10*^*-3*^, Reward Day 1: *p* = *0*.*033*, Reward Day 7: *p < 1*×*10*^*-3*^, n = 316 cells from 6 mice n = 189 cells from 7 mice **(H)**. Error bars show SEM.

To examine task-related activity in each cell type, we first compared the mean percent of active cells during the CS and reward to a null distribution made by randomly sampling the session irrespective of the behavioural task, and then calculating the mean percentage of active neurons during the sampled period. By repeating this 1,000 times for each cell type on day 1 and day 7, we created a distribution of the percentage of active neurons that were present at baseline levels or by chance. Surprisingly, we found that only PV-IN and VIP-IN cell types had a percent of CS- and reward-responsive cells that were significantly greater than chance level on both day 1 and day 7 (PV-IN CS: Day 1: 15.26 ± 2.11%, Day 7: 24.09 ± 2.98%; PV-IN reward: Day 1: 20.17 ± 3.27%, Day 7: 27.47 ± 3.66%; VIP-IN CS: Day 1: 11.29 ± 3.23%, Day 7: 16.59 ± 2.01%, VIP-IN reward: Day 1: 18.65 ± 5.91%, Day 7: 26.3 ± 6.04%; **Figure 2F, G**). PN responses to the CS were not different than the null distribution on both day 1 and 7 (Day 1: 18.42 ± 1.05%, Day 7: 16.41 ± 1.33%; **Figure 2E**); in contrast, PN responses to the reward were significantly higher on day 1 but significantly lower than the null distribution on day 7 (Day 1: 23.95 ± 2.49%, Day 7: 12.12 ± 0.95%; **Figure 2E**). Lastly, SOM-INs showed significant responses to the CS and reward only on day 7 following associative learning, while on day 1, they demonstrated no response to the CS and a modest response to the reward (CS: Day 1: 5.53 ± 2.7%, Day 7: 13.56 ± 3.17%; Reward: Day 1: 9 ± 3.66%, Day 7: 12.15 ± 3.93%; **Figure 2H**). Based on these findings, we decided to only examine the sessions where the percent of active cells were significantly greater than the null distribution in our subsequent analysis, as non-significant percent of active cells during the stimulus period cannot be readily distinguished from non-task related baseline noise.

We began our analysis on PV-INs and VIP-INs because they both showed significant responses to both CS and reward on day 1 and day 7. To understand how their representations of reward and reward-associated cues changed over the course of learning, we first analyzed the tuning of individual cells to unbiasedly identify their response properties. By quantifying the tuning of each cell’s average response during the CS and reward response periods (2.5s window) using the non-parametric Spearman correlation *ρ* (see Methods), we observed a wide range of tuning coefficients to the CS and reward, with a small proportion that was strongly positively or negatively tuned to the CS or reward stimulus (tuning coefficient near −1 or 1; **Figure 3A-D**), consistent with our earlier analyses demonstrating that neurons in M1 show activity associated with the CS or reward during the conditioning task. We next examined whether the tuning coefficient changes within each cell type after associative learning by calculating the changes in tuning coefficients for each cell between day 1 and day 7. Again, to validate our findings, we compared these values to a null distribution of *Δρ* values obtained by randomly sampling the two sessions (see Method Details). The PV-IN population did not show any significant changes in either CS or reward tuning between day 1 and 7 (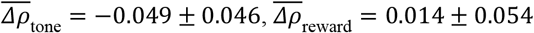; **Figure 3E, F**), indicating that neither CS-nor reward-related tuning became stronger after associative learning. In contrast, VIP-INs’ CS tuning did not change significantly between day 1 and 7 (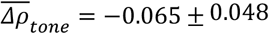, but VIP-INs’ reward tuning significantly increased on day 7 (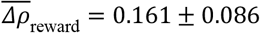; **Figure 3G, H**), suggesting a strengthening of VIP-IN responsivity to reward following associative learning.

**Figure 3.**
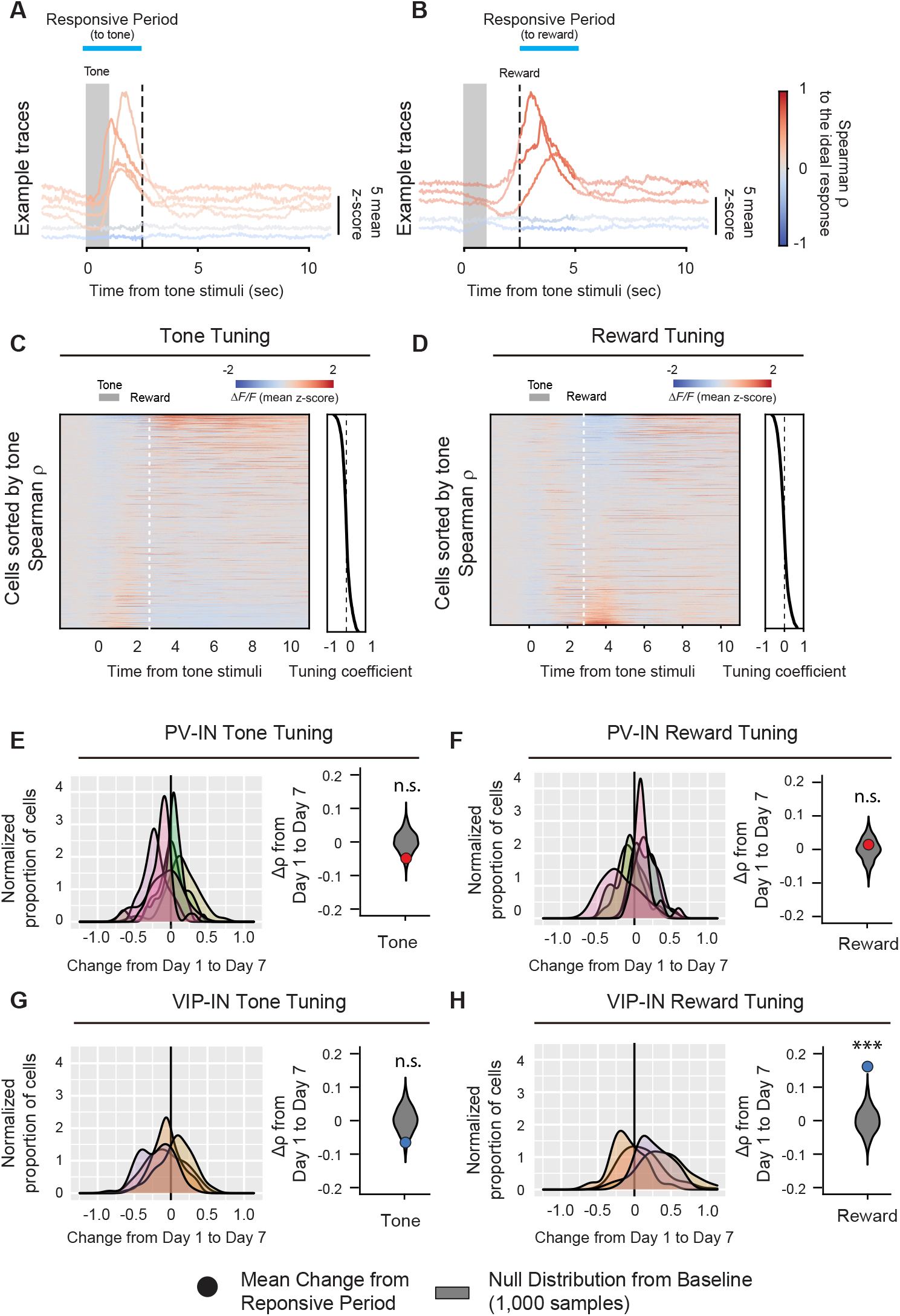
Learning-associated changes in single-neuron tuning properties in M1. (**A-B**) Example fluorescence traces, color-coded based on non-parametric Spearman correlation with tone **(A)** or reward **(B)**. Each trace is from the same neuron on day 7. (**C-D**) Trial-averaged fluorescence of all the neurons recorded on day 7 and sorted to the value of the Spearman correlation (−1 to 1) to tone **(C)** or reward **(D)**. Active neurons during the tone or reward responsive period showed higher tuning coefficient. (**E-H**) Left, distribution of changes in Spearman correlation *Δρ* with tone **(E, G)** or reward **(F, H)** for PV-INs (top) and VIP-INs (bottom). Each curve represents a Gaussian kernel density estimate of the distribution of *Δρ* in a single mouse. Right, mean change in Spearman correlation 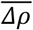 for PV-INs and VIP-INs. Null distributions (gray) was estimated by re-sampling each mouse and shuffling trials 1,000 times (see description of calculation of tuning coefficients in Methods). VIP-IN reward tuning significantly increased with associative learning. Monte-Carlo, *** *p < 1*×*10*^*-3*^, n.s., non-significant, PV-IN Tone *p = 0*.*12* **(E)**, PV-IN Reward *p = 0*.*61* **(F)**, VIP-IN Tone *p = 0*.*082* **(G)**, VIP-IN Reward *p < 1*×*10*^*-3*^ **(H)** PV-IN: n = 316 cells from 6 mice. VIP-IN: n = 407 cells from 4 mice.

Although the tuning properties can reveal changes in task-related responsivity, it is limited in identifying changes at the trial-by-trial level. When we assessed population activity following the CS onset (**Figure 4A)**, it was apparent that a group of PV-INs and VIP-INs were CS responsive on both day 1 and day 7 (**Figure 4B, 5B**). Hence, by identifying and tracking the same neurons from day 1 to day 7, we were able to ask if there was (1) an increase in the number of neurons being recruited as CS- or reward-responsive during associative learning or (2) a change in the trial-by-trial reliability of CS and reward responses. When we compared the mean percent of CS-responsive neurons on day 1 and day 7, we found that the average percent of CS responsive PV-INs during a trial increased significantly by day 7 (Day 1: 15.26 ± 2.11%, Day 7: 24.09 ± 2.98%; **Figure 4C**), while the percent of CS-responsive VIP-Ins did not change (Day 1: 11.29 ± 3.23%, Day 7: 16.59 ± 2.01%; **Figure 4D**), demonstrating that more PV-INs became responsive to the CS after associative learning. We then assessed the reliability of the responses, defined as the percent of trials within a session where a neuron was responsive to the CS. This measure quantifies how consistently a neuron responded to the CS within a session. We first plotted the cumulative distribution function of reliabilities among all PV-INs and VIP-INs. We observed that PV-INs, as a population, were significantly more reliable in their CS responses than VIP-INs on day 1 (**Figure 4E**). Next, we grouped neurons into ‘High Reliability’ if they were among the top 50^th^ percentile, while neurons in the bottom 50^th^ percentile were deemed ‘Low Reliability’. By dividing the neurons into High and Low reliability groups and following them from day 1 to day 7, we could examine if a neuron’s initial response in the naïve state will be subsequently changed by associative learning. We found that PV-INs that began as highly reliable maintained their reliability to the CS (Day 1: 29.8 ± 1.51%, Day 7: 33.87 ± 4.72%), while PV-INs that began as low reliability became significantly more reliable (8.47 ± 0.46%, Day 7: 18.99 ± 3.76%; **Figure 4F**). In contrast, the reliability of both high and low VIP-INs did not change (High Reliability: Day 1: 26.55 ± 2.62%, Day 7: 25.93 ± 3.81%; Low Reliability: Day 1: 6.32 ± 0.76%, Day 7: 14.24 ± 2.6%; **Figure 4G**). Together, these results show that as a population, more PV-INs became responsive to the CS, and their responses also became more reliable following associative learning.

**Figure 4.**
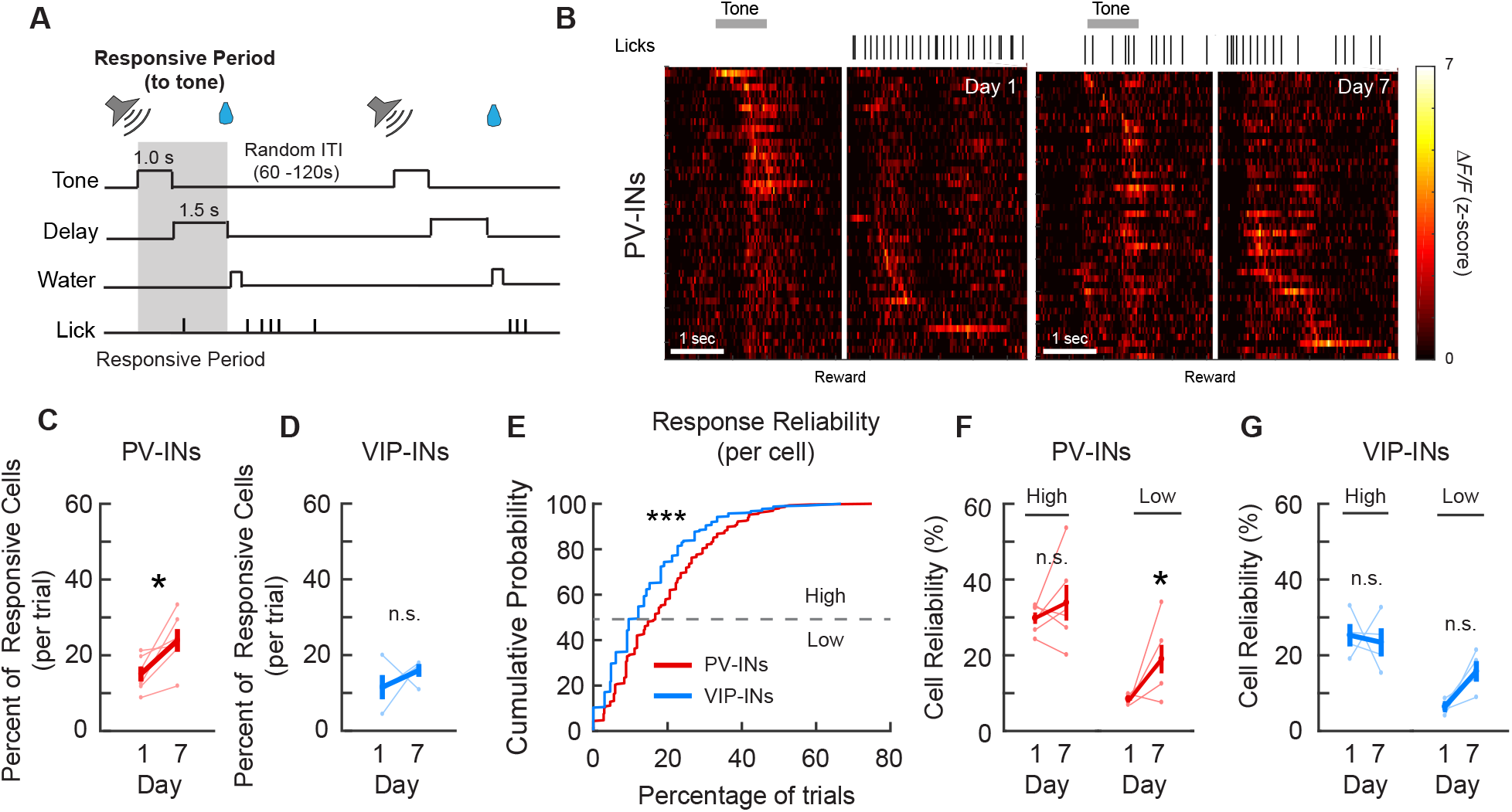
PV-IN and VIP-IN CS-related responses before and after associative learning. **(A)** Trial structure. Gray shaded bar represents the response period analyzed for tone-responsive activity. **(B)** Z-scored activity of all the active PV-INs from an example mouse during one representative trial on day 1 and day 7, sorted by timing of maximum activity following the tone onset. Grey bar represents the timing of the tone stimulus. White line indicates the onset of water reward delivery. **(C-D)** Mean percent of cells that are responsive to the tone for PV-INs (**C**) and VIP-INs (**D**). PV-INs showed an increase in the percent of tone-responsive neurons after reinforcement learning, while VIP-INs did not show any change. Paired t-test, * *p < 0*.*05*, n.s., non-significant, PV-IN: *p = 0*.*031* **(C)**, VIP-IN: *p = 0*.*38* **(D)** (**E**) Cumulative probability plots showing the percent of trials that each neuron responds to the tone for PV-INs and VIP-INs on day 1. Neurons from each cell type were pooled across mice. PV-INs showed significantly greater reliability to the tone than VIP-INs. Kolmogorov-Smirnov test, *p < 1*×*10*^*-3*^. (**F-G**) Mean reliability index of cells that are responsive to the tone for PV-INs (**F**) and VIP-INs (**G**). Each cell type is divided into High or Low Reliability Group based on the 50^th^ percentile from the cumulative probability plots in (**E**). High Reliability PV-INs maintained their consistency, while the Low Group also became more consistent in their responses to CS. **(G)** VIP-INs did not show a change in either group after reinforcement learning. Paired t-test, PV-IN high reliability: *p = 0*.*38*, PV-IN low reliability: *p = 0*.*044*, VIP-IN high reliability: *p = 0*.*79*, VIP-IN low reliability: *p = 0*.*060*. PV-IN: n = 316 cells from 6 mice. VIP-IN: n = 407 cells from 4 mice. Error bars show SEM.

We next assessed reward responses among PV-INs and VIP-INs in the same manner but now looked for responses within 2.5s of the reward delivery time (**Figure 4B, 5A-B**). We tracked the same neurons from day 1 to day 7 and compared the mean percent of reward-responsive neurons. PV-INs and VIP-INs did not show a significant change in the percent of responsive cells per trial (PV-IN: Day 1: 20.17 ± 3.27%, Day 7: 27.47 ± 3.66%; VIP-IN: Day 1: 18.65 ± 5.91%, Day 7: 26.3 ± 6.04%; **Figure 5C, D**). When we examined the cumulative distribution of reliabilities for reward responses between the two cell types, VIP-INs, as a population, were significantly more reliable than PV-INs on day 1. By dividing the cells in High and Low reliability groups, we saw that both the high and low reliability PV-INs maintained their reliability to reward (High Reliability: Day 1: 35.59 ± 2.81%, Day 7: 41.94 ± 5.7%; Low Reliability: Day 1: 10.58 ± 0.74%, Day 7: 19.59 ± 3.08%; **Figure 5F)**. In contrast, high reliability VIP-INs maintained their reliability (VIP-IN High Reliability: Day 1: 38.79 ± 2.71%, Day 7: 35.88 ± 4.32%), and the low reliability VIP-INs became significantly more reliable on day 7 (VIP-IN Low Reliability: Day 1: 10.25 ± 1.67% Day 7: 24.06 ± 4.87%; **Figure 5G**). Altogether, while the proportion of reward-responsive VIP-INs during a given trial did not change, a subset of VIP-INs that were largely unresponsive to reward on day 1 became more reliably responsive on day 7.

**Figure 5.**
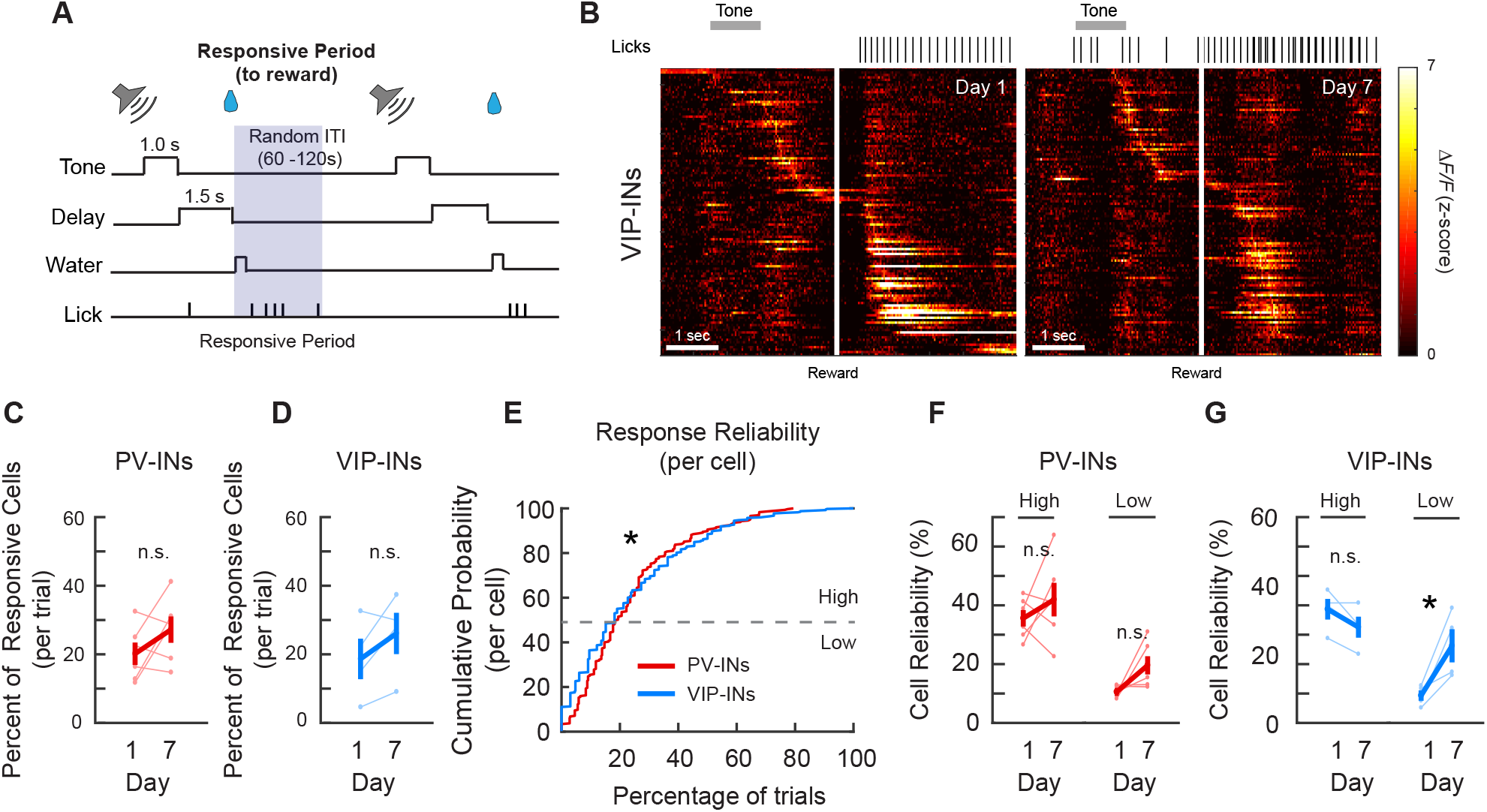
PV-IN and VIP-IN reward-related responses before and after associative learning. **(A)** Trial structure. Gray shaded bar represents the response period analyzed for reward-responsive activity. **(B)** Z-scored activity of all the active VIP-INs from an example mouse during one representative trial on day 1 and day 7, sorted by timing of maximum activity following the tone onset. Grey bar represents the timing of the tone stimulus. White line indicates the onset of water reward delivery. **(C-D)** Mean percent of cells that are responsive to the reward for PV-INs (**C**) and VIP-INs (**D**). Neither PV- or VIP-INs showed a significant change. Paired t-test, n.s., non-significant, PV-IN: *p = 0*.*16* **(C)**, VIP-IN: *p = 0*.*16* **(D)** (**E**) Cumulative probability plots showing the percent of trials that each neuron responds to reward for PV-INs and VIP-INs on day 1. Neurons from each cell type were pooled across mice. VIP-INs showed a significantly greater response reliability to the reward than PV-INs. Kolmogorov-Smirnov test, * *p < 0*.*05, p = 5*.*2* × *10*^*-3*^. (**F-G**) Mean reliability index of cells that are responsive to the reward for PV-INs (**F**) and VIP-INs (**G**). Each cell type is divided into High or Low Reliability Group based on the 50^th^ percentile from the cumulative probability plots in (**E**). High and Low Reliability PV-INs maintained their consistency. (**G**) High Reliability VIP-INs did not show a change, while Low Reliability VIP-INs significantly increased in reliability following associative learning. Paired t-test, PV-IN high reliability: *p = 0*.*36*, PV-IN low reliability: *p = 0*.*058*, VIP-IN high reliability: *p = 0*.*090*, VIP-IN low reliability: *p = 0*.*045*. PV-IN: n = 316 cells from 6 mice. VIP-IN: n = 407 cells from 4 mice. Error bars show SEM.

Although PV-INs and VIP-INs were the only cell types that were significantly responsive to both CS and reward on both day 1 and day 7, PNs and SOM-INs also had significant responses to specific stimuli on certain days. While PNs did not show significant CS responses when compared to baseline, their reward responses on day 1 were significantly above the null distribution, and they became significantly lower than the null distribution on day 7 (**Figure 2E**). This result is in line with the change in tuning coefficient (Δ*ρ*_*reward*_), which showed a significant decrease in reward tuning between day 1 and 7 (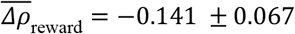; **Figure 6A**). Moreover, the cumulative distribution function of PN reliability also shifted significantly to lower reliabilities on day 7 compared to day 1 (**Figure 6B**). These results indicate that PNs initially respond to novel reward; however, they habituate to the reward following associative learning.

**Figure 6.**
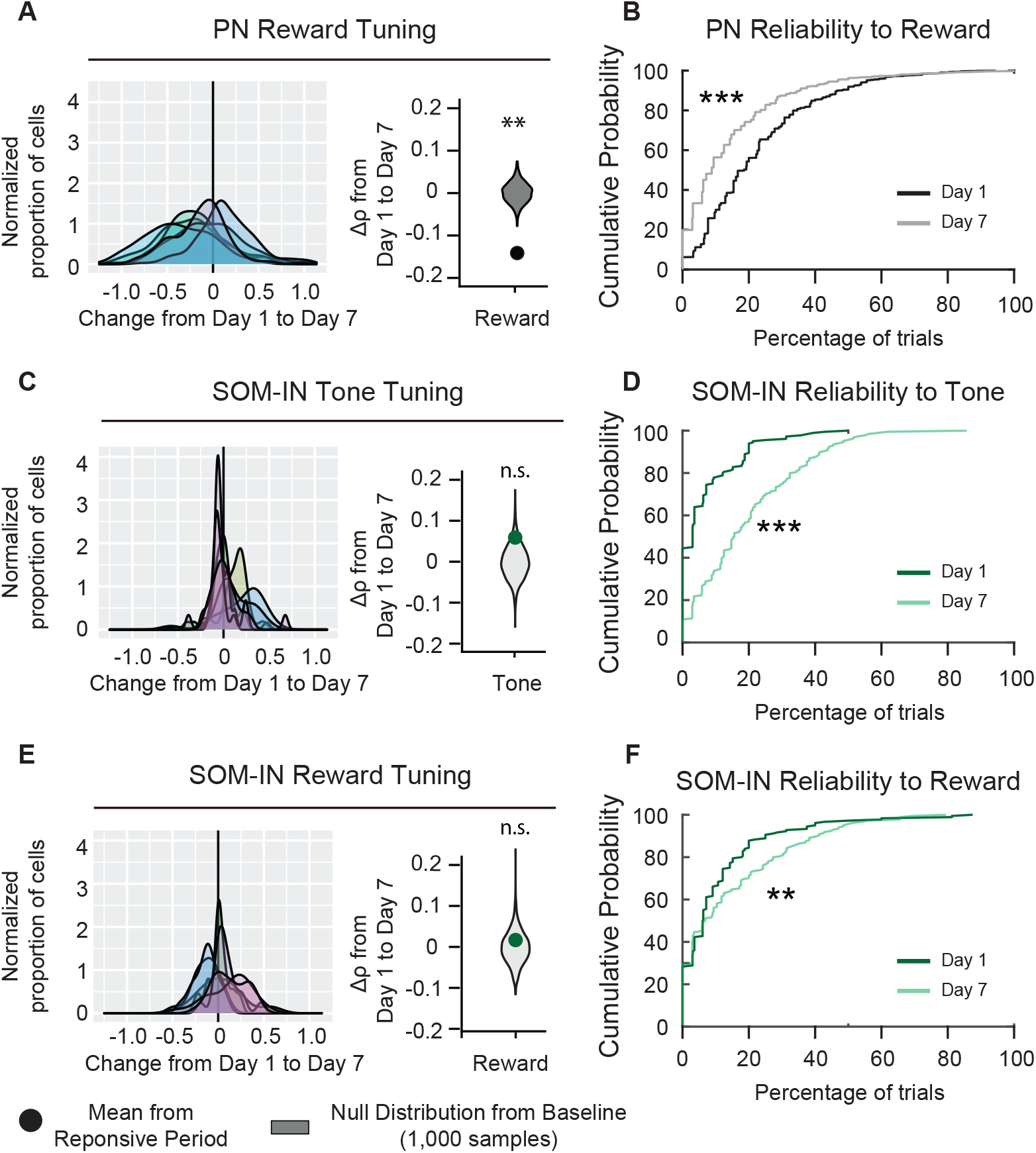
PN and SOM-IN reliability is altered after associative learning. **(A)** Left, distribution of changes in PN Spearman correlation *Δρ* for reward. Each curve represents a Gaussian kernel density estimate of the distribution of 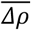 in a single mouse. Right, mean change in Spearman correlation *Δρ*. Null distributions (gray) is estimated by re-sampling each mouse and shuffling trials 1,000 times. Reward tuning among PNs decreased after associative learning, Monte Carlo, *** *p < 1*×*10*^*-3*^. **(B)** Cumulative probability plots showing the percent of trials that each neuron responds to reward for PNs on day 1 and day 7. Neurons were pooled across mice. Day 7 reliability was significantly lower than day 1. Kolmogorov-Smirnov test, *** *p < 1*×*10*^*-3*^. **(C)** Left, distribution of changes in SOM-IN Spearman correlation *Δρ* with tone. Each curve represents a Gaussian kernel density estimate of the distribution of *Δρ* in a single mouse. Right, mean change in Spearman correlation 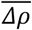. SOM-INs did not show a change in tone tuning. Monte Carlo, n.s., non-significant, *p = 0*.*128*. **(D)** Cumulative probability plots showing the percent of trials that each neuron responds to the CS for SOM-INs on day 1 and day 7. Neurons were pooled across mice. Day 7 reliability was significantly greater than day 1. Kolmogorov-Smirnov test, *p < 1*×*10*^*-3*^. **(E)** Left, distribution of changes in SOM-IN Spearman correlation *Δρ* with reward. Each curve represents a Gaussian kernel density estimate of the distribution of *Δρ* in a single mouse. Right, mean change in Spearman correlation 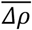. SOM-INs did not show a change in reward tuning. Monte Carlo, n.s., non-significant, *p = 0*.*598*. **(F)** Cumulative probability plots showing the percent of trials that each neuron responds to the reward for SOM-INs on day 1 and day 7. Neurons were pooled across mice. Day 7 reliability was significantly greater than day 1. Kolmogorov-Smirnov test, ** *p = 0*.*012*. PN: n = 1029 cells from 6 mice. SOM-INs: n = 189 cells from 7 mice.

SOM-INs initially had no response to the CS on day 1, but their responses became significant on day 7 (**Figure 2H**). The change in CS tuning coefficient (Δ*ρ*_*tone*_) was not significant (Δ*ρ*_*tone*_ = .059 ± 0.031, **Figure 6C**), suggesting their responsivity did not change with learning. Interestingly, when we assessed the change in reliability to the CS between day 1 and day 7, the cumulative distribution function shifted significantly to higher reliability values on day 7 (**Figure 6C, D**). Notably, by day 7, there was a visible reduction in the number of SOM-INs that had 0% reliability to CS on day 1, indicating they were completely unresponsive to tone on day 1 but not on day 7. Finally, SOM-INs showed modest but significant responses to reward on day 1 and 7 (**Figure 2H**). When we assessed the reward tuning among the SOM-IN population, Δ*ρ*_*reward*_ did not show a significant change between day 1 and 7 (Δ*ρ*_*reward*_ = .0.017 ± 0.040, **Figure 6E**). However, SOM-IN reliability also shifted to higher values on day 7 (**Figure 6F**). Altogether, these results suggest that in naïve mice, SOM-INs are unresponsive to the neutral CS and modestly responsive to the novel reward stimulus; however, following associative learning, SOM-INs become more reliably responsive to both the CS and reward.

Lastly, reward consumption requires innate tongue movements during licking, and since microstimulation of mouse M1 has been shown to evoke tongue and jaw movements (Komiyama et al., 2010), it is crucial to distinguish whether the observed CS and reward responses resulted from the task-related stimuli or if the activity is simply associated with licking movements. We demonstrated earlier that head-fixed mice learned the CS-US association by displaying the conditioned response (anticipatory licking) following the CS on day 7 (**Figure 1**). To address this potential confound, we identified all the self-initiated licking bouts during ITIs, when no reward was present (**Figure 7A-C)**. We assessed all the significantly active cell types on day 1 and day 7 (identified in Figure 2) and calculated the response reliability index of all the active neurons during ITI lick bouts and compared them to the response reliability index for the CS and reward. On both day 1 and day 7, all cell types exhibited lower reliability index values for the ITI lick bouts compared to the CS and reward, indicating that the increase in task-related responses following water rewards was specific to the reward stimulus, and not licking movements (**Figure 7D-J)**. These results suggest that the cell-type specific modifications observed between day 1 and 7 were not simply caused by licking movements.

**Figure 7.**
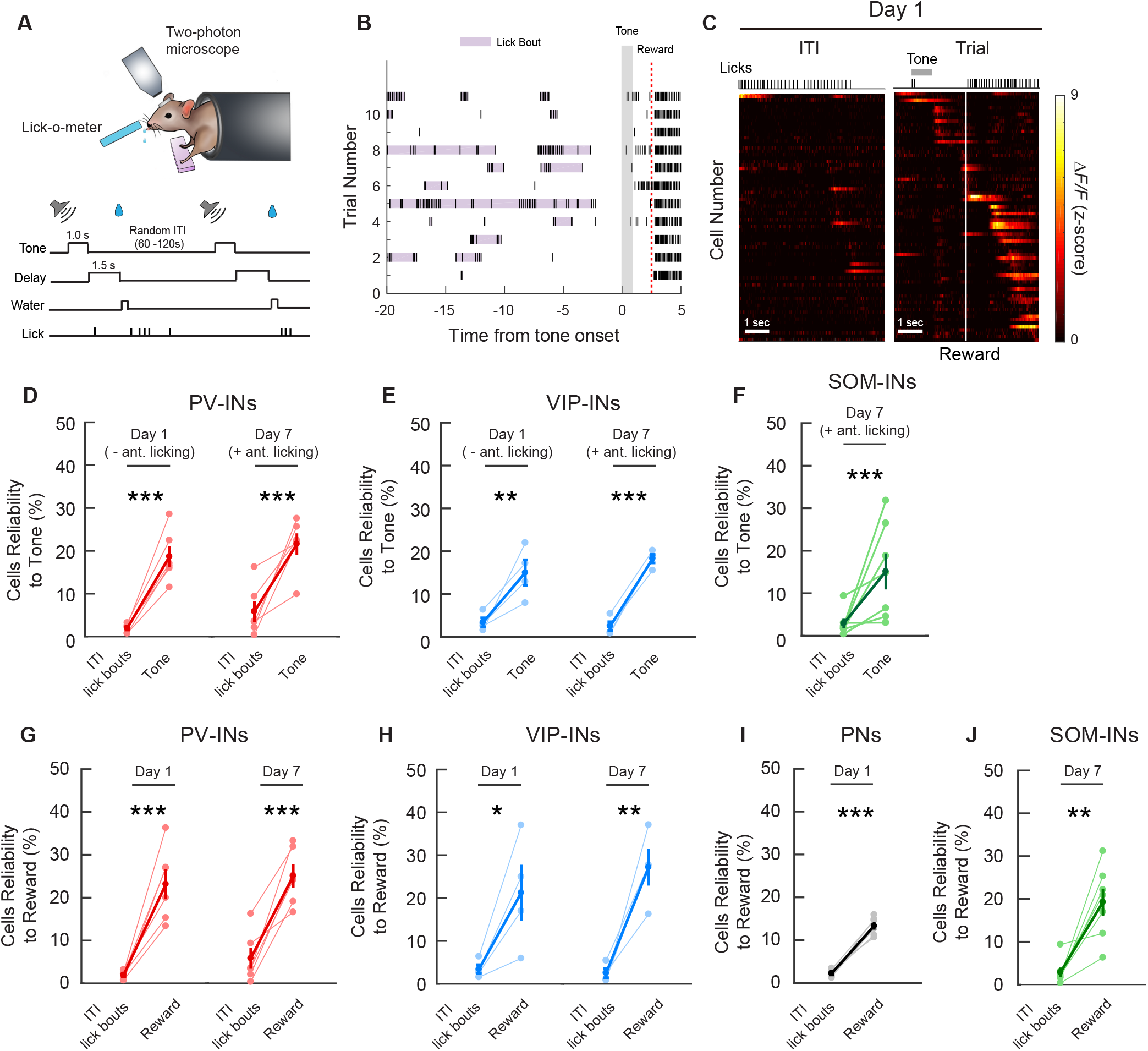
Cell-type specific cue and reward activity are not due to licking movements. **(A)** Schematic of the cued reward conditioning task (top) and the trial structure (bottom). **(B)** Example licking behaviour during the ITIs from one mouse on day 1. Purple shading shows licks that were considered to be an individual lick bout. Gray shaded bar shows the tone timing and the red dotted line shows the reward timing. **(C)** Z-scored activity of all the active PNs from an example mouse during one representative ITI lick bout. Left: maximum activity aligned to the lick bout onset. Right: maximum activity aligned to the tone onset. Grey bar represents the timing of the tone stimulus. White line indicates the time of water reward delivery. **(D-F)** Mean reliability index for all cell types with significant tone-related responses during ITI lick bouts with no water reward present, compared to the mean reliability index during the tone and up to but not including the reward delivery time. PV-INs, VIP-INs and SOM-INs were more reliably responsive during the CS than during licking movement alone. Only sessions/cell-types with significant CS-related responses were analyzed. Paired t-test, * *p < 0*.*05*, ** *p < 0*.*01*, *** *p < 0*.*001*. n.s., non-significant. PV-INs Day 1: *p < 1*×*10*^*-3*^, PV-IN Day 7: *p = 6*.*6*×*10*^*-3*^ **(D)**, VIP-IN day 1: *p = 0*.*0355*, VIP-IN day 7: *p < 1*×*10*^*-3*^ **(E)**, *SOM-IN Day 7: p = 3*.*9*×*10*^*-3*^ **(F)**. **(G-I)** Mean reliability index for all cell types with significant reward-related responses during ITI lick bouts with no water reward present, compared to the mean reliability index following reward timing. All cell types were more reliably responsive during the reward period than during licking movement alone. Only sessions/cell-types with significant reward-related responses were analyzed. Paired t-test, PV-INs Day 1: *p =1*.*6*×*10*^*-3*^, PV-IN Day 7: *p = 1*.*6* × *10*^*-3*^ **(G)**, VIP-IN day 1: *p = 0*.*049*, VIP-IN day 7: *p = 0*.*014* **(H)**, PN Day 1 *p < 1*×*10*^*-3*^ **(I)**, SOM-IN Day 7: *p = 0*.*030* **(J)**. PN: n = 1029 cells from 6 mice. PV-IN: n = 316 cells from 6 mice. VIP-IN: n = 407 cells from 4 mice. SOM-IN: n = 189 cells from 7. Error bars show SEM.

## DISCUSSION

M1 is known to be involved in motor initiation, movement kinematics and motor learning. Recent studies have demonstrated reward-related activity in M1 using *in vivo* electrophysiological recordings in nonhuman primates (Marsh et al., 2015; Ramakrishnan et al., 2017; Ramkumar et al., 2016) and transcranial magnetic stimulation in human subjects (Thabit et al., 2011). However, whether CS- and reward-associated signals are represented among different neuronal cell types within the microcircuit in M1 is still unclear. Using chronic two-photon Ca^2+^ imaging, combined with transgenic mouse lines and viral strategies to target different neuronal cell types, we demonstrated that during a conditioning task, all major cell types in M1 responded to either the CS, the reward stimulus, or both. Most notably, each cell type underwent cell-type specific modifications after association learning. By tracking the same population of neurons before and after associative learning, we revealed that the CS-responding population increased among PV-INs, and individual cell responses to the CS also became more reliable following associative learning. On the contrary, VIP-INs became more reliable to reward. Additionally, PNs had a drastically reduced response to reward, while SOM-INs became more reliable to both the CS and the reward. Our findings suggest that each cell type has a distinct role in processing information related to the cue-reward association in M1, and they may work together to provide the reinforcement signals in M1 that are important for motor skill learning.

Previous studies in trained rhesus monkeys performing a joystick center-out task have shown a widespread representation of reward anticipation and reward-related activity among cortical neurons in M1 (Ramakrishnan et al., 2017). Consistent with earlier work, we also observed reward-related activity in all four major cell types in M1, even in naïve mice on day 1 when they were first exposed to the CS and reward. It has been reported that in sensory cortices, repeated passive exposure to a sensory stimulus leads to a long-lasting reduction in PN responsivity, but when animals are engaged in learning, PNs maintain their responsivity to the repeated stimulus (Kato et al., 2015; Makino & Komiyama, 2015). However, we found in M1, when water-restricted mice were engaged in a conditioning task to learn the association between the CS and water reward, PNs still showed a drastic habituation to the reward stimulus. A recent study that imaged neuronal activity in expert mice performing a head-fixed pellet reaching task demonstrated that L2/3 PNs in M1 are involved in encoding movement outcome (success vs. failure) but not the appetitive outcome (reward vs. no reward). However, the authors did not image the mice at the naïve stage (Levy et al., 2020). Hence, one possibility is that L2/3 PNs in M1 encode reward signals during the naïve stage, but after associative learning, they habituate and become unresponsive to the reward stimulus. In addition, in the sensory cortices, the flexibility to either respond to or ignore sensory stimuli is based on the stimulus’ behavioral relevance, and it is gated by local SOM-INs (Kato et al., 2015; Makino & Komiyama, 2015). In line with these findings, we found that SOM-INs became more reliably responsive to both the CS and the reward stimulus with associative learning. We also observed stimulus-specific increases in reliability in PV-INs’ response to the CS and VIP-INs’ response to the reward after associative learning, suggesting that different IN subtypes may have distinct roles in processing CS- and reward-related information in M1.

One hypothesis is that PV-INs are recruited by the CS to regulate anticipatory licking during reward anticipation since PV-INs are known to regulate PN firing through both feedforward and feedback inhibition (Fishell & Rudy, 2011; Xu & Callaway, 2009; Xue et al., 2014). Similar observations have been reported in the striatum, in which optogenetic activation or suppression of PV-INs during a similar conditioning task impaired anticipatory licking, demonstrating the importance of PV-INs in the expression of conditioned responses (Lee et al., 2017). Likewise, PV-INs in the basolateral amygdala, are also recruited during the CS and subsequently inhibited during the US in an auditory fear conditioning task. Optogenetic activation of PV-INs during the CS increased conditioned freezing behaviour while PV-IN suppression reduced freezing, indicating bidirectional control of the conditioned response (Wolff et al., 2014). Our results demonstrate that in M1, a subset of PV-INs was already highly responsive to the CS before conditioning (day 1, naïve stage), and more PV-INs were recruited by the CS after associative learning, suggesting PV-INs in M1 might also play an important role in controlling conditioned responses.

VIP-INs, on the other hand, were significantly less reliable in responding to the CS compared to PV-INs, and their responses to the CS remained low. However, VIP-INs’ responses to the reward were more reliable than those of PV-INs, and they became more closely tuned and reliably responsive to the reward with learning. Due to the disinhibitory position of VIP-INs in the microcircuit, activation of VIP-INs can lead to widespread increases in local excitability and contribute to cortical gain control (Fu et al., 2014; Jackson et al., 2016; Pfeffer et al., 2013). Furthermore, a growing body of evidence suggests a general principle across brain regions, in which VIP-INs receive long-range inputs (Duan et al., 2020; Krabbe et al., 2019; Turi et al., 2019; Zhang et al., 2014), respond to reinforcement signals (Krabbe et al., 2019; Pi et al., 2013), and play an important role in goal-oriented learning (Krabbe et al., 2019; Turi et al., 2019). Taken together, our results suggest that during CS-reward conditioning, PV-INs in M1 are critical for encoding the CS association and regulate local circuit activity related to reward anticipation, whereas VIP-INs act as a context-dependent switch following the reward delivery (Muñoz et al., 2017; Turi et al., 2019) to instruct and disinhibit local PNs to enable learning-induced plastic changes critical for the acquisition of new movements. This work may provide insight on how different IN subtypes in M1 integrate incoming inputs from various brain regions and orchestrate local circuit plasticity.

## ACKNOWLEDGEMENTS

We thank the members of the Chen lab for discussions and providing feedback on the manuscript. This work was supported by grants for S.X.C. from Canada Research Chair (CRC) (Grant no. 950-231274) and Natural Sciences and Engineering Research Council of Canada (NSERC) (Grant no. 05308), and a grant for R. N. from NSERC (Grant no. 06972). E.H. was supported by a NSERC graduate scholarship. C.L. was supported by Ontario Graduate Scholarship and Queen Elizabeth II Graduate Scholarship.

## COMPETING INTERESTS

The authors declare no competing interests.

## AUTHOR CONTRIBUTIONS

C.L. and S.X.C. conceived the project. C.L. conducted all the experiments. E.H. and R.N. performed the analyses for Figure 3. Rest of the figures were analyzed by C.L. under supervision of S.X.C. C.L. and S.X.C. wrote the manuscript.

## MATERIALS AND METHODS

All animal experiments were approved by the University of Ottawa Animal Care Committee and in accordance with the Canadian Council on Animal Care guidelines. Experimental mice were group-housed in plastic cages with food and water ad libitum in a room with a reversed light cycle (12h - 12h). Mice were acquired from Jackson Laboratory (Bar Harbor, ME, USA; PV-Cre (008069), SOM-Cre (013044), VIP-Cre (010908), B6129SF1/J (101043)). For all mouse lines, both male and females were used. Mice were between P40 and P60 at the time of surgery.

### Surgery

Mice were deeply anesthetized under 1-2% isoflurane and given subcutaneous injections of Batyril (10 mg/kg) to prevent infection and buprenorphine (0.05 mg/kg) for analgesia. An incision was performed to remove a piece of the scalp and a custom head-plate was implanted onto the skull using instant glue (Krazy Glue) and dental cement (Lang Dental, Wheeling, IL, USA). A craniotomy of approximately 2 mm in diameter was performed over the right primary motor cortex. Virus (PNs: AAV1.CaMKII.GCaMP6f.WPRE.SV40; PV-IN, VIP-IN and SOM-IN: AAV1.Syn.Flex.GCaMP6f.WPRE.5v40) was diluted 1:5 in saline and injected at a depth of ∼250 µm from the pia using a glass pipette. All virus was obtained from Addgene (Watertown, MA, USA). Injections were performed at 5 sites, centered on coordinates 1.5 mm lateral and 0.3 mm anterior to bregma. For PN groups, 20 nl per site was injected. For PV-IN, VIP-IN and SOM-IN, 40 nl per site was injected. All injections were performed at a rate of 10 nl/min and the pipette was left in place for 4 minutes following the injection to avoid backflow. A glass imaging window was then implanted over the craniotomy and sealed with dental cement. Following surgery, a subcutaneous injection of dexamethasone (2 mg/kg) and buprenorphine (0.1 mg/kg) was given. Mice were given a minimum of 1 week to recover prior to beginning water restriction.

### Auditory Cued Reward Conditioning Behaviour

Mice were gradually water restricted down to ∼1 ml per day (∼80% of original body weight) over two weeks prior to the start of imaging/behaviour sessions (Chen et al., 2015; Harvey, Coen, & Tank, 2012; Komiyama et al., 2010; O’Connor et al., 2013; Peters et al., 2014). Mice were then head-fixed for simultaneous two-photon imaging and exposed to the unconditioned stimulus (a constant auditory tone, 1s in duration) followed by a 1.5s delay period and a water reward (∼ 10 µl). All lick times were measured by an infrared beam lick-o-meter and logged using the data acquisition software WaveSurfer (https://wavesurfer.janelia.org/). The inter-trial interval between the previous water reward and subsequent tone onset was randomly varied between 60-120s. Each session was one hour in duration with 30-35 trials in total. Mice underwent one session per day for seven consecutive days. Two-photon calcium image was performed simultaneously on day 1 and day 7 of the behavioural task.

To assess licking behaviour, lick rate (number of licks per second, measured as infrared beam breaks) was calculated within 500ms bins, then averaged across all trials within a session for each mouse. Lick rate was then averaged across mice. Mean anticipatory lick rate was calculated as the mean lick rate from the time of tone onset to the end of the delay period (2.5s in duration), not including the reward delivery. Mean ITI lick rate was calculated from the lick rate during the first 2.5s of self-initiated spontaneous lick bouts. ITI lick bouts were defined as licking events that followed the previous trial by at least 20s and preceded the subsequent trial by more than 2.5s. Mean reward lick rate was calculated from the lick rate from the time of reward delivery to 2.5s after.

All trials within a session were including in lick rate analysis in Figure 1. To ensure behavioural consistency across trials, only trials with at least 3 lick responses within 2.5 s of the reward delivery time were included in all analysis of neural responses.

### Calcium Imaging and Analysis

*In vivo* imaging was performed using a commercial two-photon microscope (B-scope, Thorlabs, Newton, NJ, USA) and a 16x water immersion objective (Nikon) with excitation at 925 nm (InSight X3, Spectra-Physics, Milpitas, CA, USA) with a frame rate of 30 Hz. Images were taken at 512 × 512 pixels covering 755 by 650 µm.

Images were corrected for movement in the x and y plane using full-frame cross-correlation image alignment (Turboreg (Thévenaz, Ruttimann, & Unser, 1998) plug-in ImageJ). The entire session was visually inspected and regions of interests (ROIs) were manually drawn on neurons using a custom MATLAB program, described in Peters et al. (Peters et al., 2014). The ROI template from day 1 was loaded onto day 7 and aligned along the x and y plane. Only neurons that could be tracked from day 1 to day 7 were included in the dataset.

Fluorescence within an ROI was averaged across pixels. ΔF was calculated by subtracting the baseline fluorescence (F_0_) from the raw fluorescence trace. The calculation for baseline fluorescence (F_0_) was based on inactive parts of the fluorescence trace as previously described (Chu, Li, & Komiyama, 2016; Kato, Chu, Isaacson, & Komiyama, 2012; Peters et al., 2014; Peters, Lee, Hedrick, O’Neil, & Komiyama, 2017)

We adapted a method by Driscoll et al. (Driscoll, Pettit, Minderer, Chettih, & Harvey, 2017) to identify significant activity events for each neuron and then excluded ROIs with no significant activity events within the session, irrespective of the behaviour. For each neuron the ΔF trace was circularly shifted by a random integer 1,000 times and compared to the original trace. If the original ΔF trace was greater than the shifted data for at least 5 consecutive frames in at least 950 iterations, this was considered an active event. If a neuron did not have at least one active event in the entire session, irrespective of the behaviour, it was removed from the data set. This only accounted for a small proportion of ROIs as most of them are active on both day 1 and day 7, as shown in Figure 2D.

For all subsequent analyses, a modified z-score, adapted from Kato et al. (2015), was applied to ΔF. The z-score was calculated as *Z = (f(t) - µ)/σ*, where *f(t)* is the ΔF trace for a neuron, *µ* is the mean, and *σ* is the standard deviation of the neuron’s ΔF during the baseline period. The baseline period was a concatenation of 2.5 s preceding the tone onset (start of a trial) for all trials within a session.

### Calculation of Tuning Coefficients

We quantified the tuning of individual neurons to the tone and reward stimuli delivered in our classical conditioning task using the non-parametric Spearman correlation *ρ* (scipy.stats.spearmanr) between the trial-averaged fluorescence and the timing of stimulus delivery

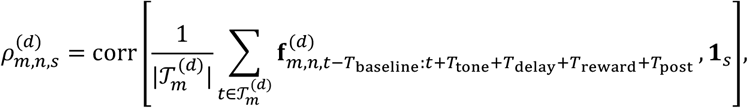

where *t* is the start time of a trial (defined as the start of the tone stimulus), 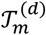 is the set of all trial start times from mouse *m* on day *d* ∈ {1,7}, 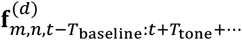 is the fluorescence trace of neuron *n* from mouse *m* during a single trial, **1**_*s*_ is an indicator function for stimulus *s* ∈ {tone, reward}, 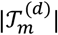 is the number of trials, and *ρ* is the Spearman correlation coefficient. Analysis was carried out with *T*_baseline_ =2 s and *T*_post_ =6 s. We considered the “tone” period indicated by **1**_tone_ to range from the start of the tone at time *t* to the start of reward delivery at time *t* + *T*_tone_ + *T*_delay_, and the “reward” period indicated by **1**_reward_ to be the first 2.5 seconds of reward delivery (see schematic in Figure 4A, 5A). We used the change in *ρ* from day 1 to day 7 as a cell-resolved measure of changes in tuning over the course of learning.

To summarize learning-associated changes in tuning, we calculated the mean change in the Spearman correlation for each cell type and trial component (tone or reward) from day 1 to day 7 as follows

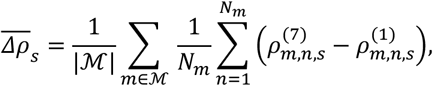

where ℳ is the set of mice used in the experiment, *N*_m_ is the number of neurons in mouse *m*, and 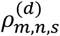 is the Spearman correlation as defined above.

We used a non-parametric approach for statistical tests involving the mean change in Spearman correlation by scrambling trial times and bootstrapping mice to construct a null distribution for 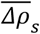. Specifically, we first drew a random sample of |ℳ| mice from ℳ with replacement, then drew a random sample of 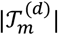 trial start times uniformly distributed between 0 and 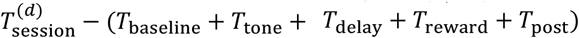 for each day *d* and randomly-selected mouse, and finally used these randomly-selected mice and scrambled trial start times to compute the change in tuning 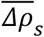. This process was repeated 1000 times to approximate the distribution of 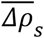 under the null hypothesis that changes in tuning are unrelated to tone and reward delivery. We considered the observed changes in tuning 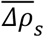 to be statistically significant at the ∗ or ∗∗ level if they fell into the 5 % or 1 % tails of this distribution, respectively.

### Activity Analysis

To identify neuron responses to tone and reward, we applied a set threshold to each neuron on a trial-by-trial basis. Neurons were defined as tone-responsive or reward-responsive within a trial if they exceeded 1 z-score (excitation threshold used in Kato et al., 2015 (Kato et al., 2015)) for at least 5 consecutive frames within 2.5s of the tone onset or 2.5s of the reward delivery time, respectively. This was assessed for each trial with at least 3 lick responses within 2.5 s of the reward delivery time. We then took the median of the percent of responsive neurons across all trials in a session from one mouse, and the mean across mice.

We used a Monte-Carlo approach to validate the percent of tone and reward responsive neurons. The mean percentage of tone-responsive and reward-responsive neurons observed were compared to a null distribution made for each cell type on each day. We randomly sampled mice with replacement, then sampled the entire session, and then calculated the percentage of active cells (exceeding 1 z-score for at least 5 consecutive frames) during a randomly chosen 2.5s window. For each mouse, the number of samples was equal to the number of included trials (i.e. number of trials with at least 3 lick responses within 2.5s of reward delivery). We then took the median across the random samples and then took the mean across mice to obtain a mean percentage of responsive neurons during a randomly chosen time window. This was repeated 1000 times to generate a null distribution of mean percentage of active neurons. A Monte-Carlo approach was then used to assess whether the observed percentage of tone- and reward-responsive neurons was significantly different from the null distribution by comparing the observed value to the tails of the normally distributed null distribution. This was done for each cell type on both Day 1 and Day 7. We considered the tone- or reward-responses to be statistically significant at the ∗ or ∗∗ level if they fell into the 5 % or 1 % tails of this distribution, respectively and *** if there was no overlap with the distribution. Since this approach tests the null hypothesis that the observed neuronal responses are due to chance (in this case, baseline activity/noise), only cell types with a significantly higher percentage of responsive neurons for a given session were analyzed.

The tone/reward reliability index was defined as the percent of trials within a session where the neuron was tone/reward responsive. The reliability cumulative distribution was made by pooling the day 1 index values of all the neurons from a neuronal cell type (across mice). If a neuron’s day 1 index value was lower or equal to the index value at the 50^th^ percentile of the cumulative distribution for that cell type, it was categorized into the Low Reliability group. If a neuron’s day 1 index value exceeded the 50^th^ percentile value, it was categorized into the High Reliability group. We then took the mean reliability within each group on day 1 and day 7.

### Inter-Trial Interval Lick Bout Analysis

Inter-trial interval (ITI) lick bouts were defined as self-initiated licking events that occurred at least 20 s after the preceding reward delivery time (trial end) and more than 2.5s prior to the subsequent tone onset (trial start). If licks were separated by 3s or more, they were considered a new lick bout. To remain consistent with tone and reward analyses, only the first 2.5s of a lick bout were analyzed for neural responses. ITI lick bout reliability indices were calculated as described above.

### Statistical Analysis

Statistical analysis for tuning coefficients were performed in Python and in R. Statistical analyses for anticipatory licking, tone- and reward-responsivity, and reliability index were performed in Matlab using the Statistics and Machine Learning Toolbox. Two-way ANOVA was used to test for differences in anticipatory lick rate on day 1 and day 7. One-way ANOVA was used to test for differences in lick rate during ITI, tone and reward. One-way ANOVA was used to compare the percent of active cells across cell types on a single day. Monte-Carlo (as described above) was used to test for significant percent of tone- and reward-responsive neurons, and for changes in tuning properties. Paired t-test was used to test for differences in the percentage of responsive cells and reliability index on day 1 and 7, and for differences in neuron reliability between ITI lick bouts, tone and reward. The Kolmogorov–Smirnov test was used to compare response reliability cumulative distributions. All values were reported as the mean and standard error of the mean unless otherwise specified. Power analysis was not performed to predetermine the sample size, and the experiments were not blinded.

### Data Analysis and Code Availability

Tuning coefficient calculation and statistical tests were performed using Python 3.8 with the following libraries: NumPy, Pandas, h5py, and SQLAlchemy. Figures were prepared in Python using matplotlib and seaborn, and in R using ggplot2. Codes to reproduce the analysis for figures 1-2 and 4-7 are available at https://github.com/clee162/Analysis-of-Cell-type-Specific-Responses-to-Associative-Learning-in-M1. Codes to reproduce the analysis and figure 3 are available at https://github.com/nauralcodinglab/interneuron-reward. Data can be found on Dryad at https://doi.org/10.5061/dryad.q573n5tjj.

## Notes

### Competing Interest Statement

The authors have declared no competing interest.

